## Supplementary figures and images for "Bidirectional fear modulation by discrete anterior insular circuits in male mice"

### Figure 3-figure supplement 1

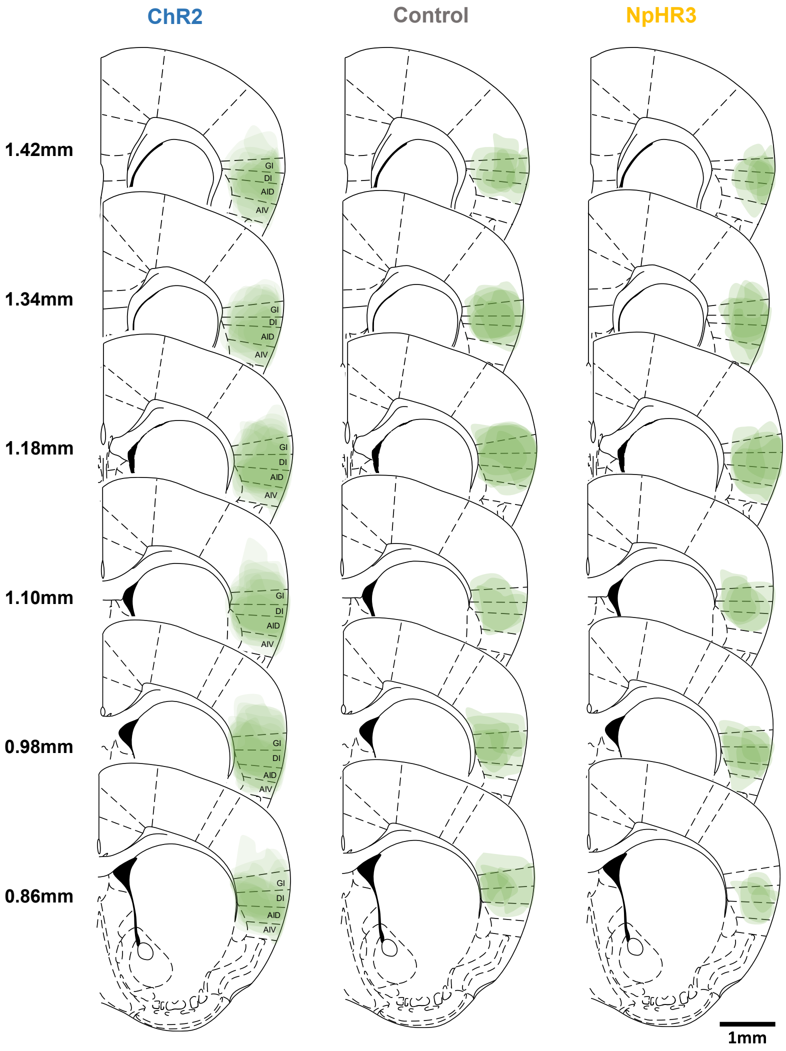

### Figure 4-figure supplement 1

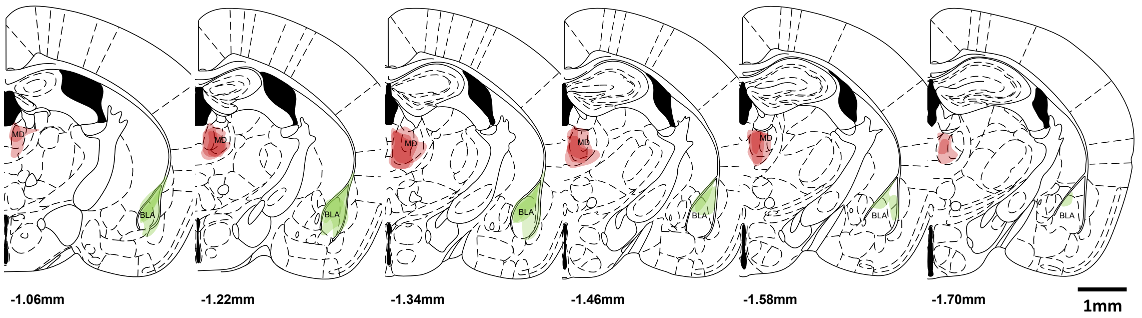

### Figure 5-figure supplement 1

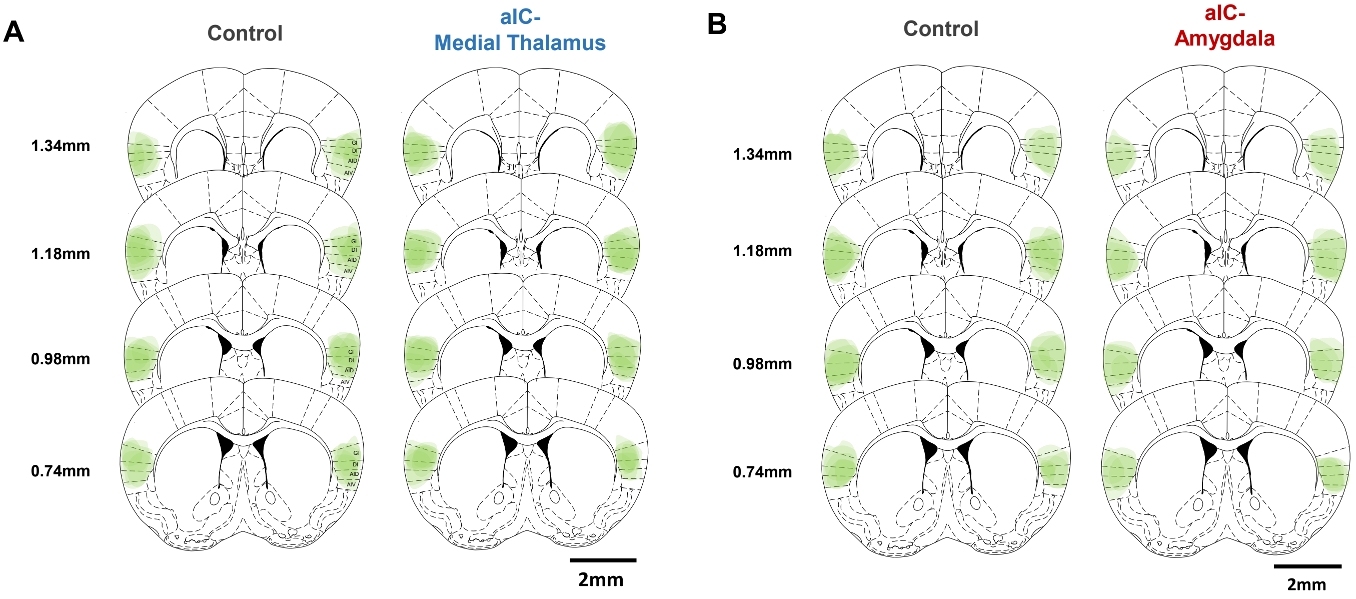
